# *Methanolobus* use unspecific methyltransferases to produce methane from dimethylsulfide

**DOI:** 10.1101/2023.08.04.551946

**Authors:** S. L. Tsola, Y. Zhu, Y. Chen, I. A. Sanders, C. K. Economou, V. Brüchert, Ö. Eyice

**Affiliations:** School of Biological and Behavioural Sciences, Queen Mary University of London, London, UK; School of Life Sciences, University of Warwick, Coventry, UK; Department of Geological Sciences, Stockholm University, Stockholm, Sweden; Bolin Centre for Climate Research, Stockholm University, Stockholm, Sweden

## Abstract

Dimethylsulfide (DMS) is the most abundant biogenic organic sulfur compound and a methane precursor in anoxic sediments. However, understanding of the microbial diversity driving DMS-dependent methanogenesis is limited, and the metabolic pathways underlying this process in the environment remain unexplored. To address this, we used anoxic incubations, amplicon sequencing, genome-centric metagenomics and metatranscriptomics of brackish sediments of the Baltic Sea. We identified *Methanolobus* as the dominant methylotrophic methanogens in all our sediment samples. We also showed that *Methanolobus* use trimethylamine- and methanol-methyltransferases, not methyl-sulfide methyltransferases, when producing methane from DMS. This demonstrated that methylotrophic methanogenesis does not require a substrate-specific methyltransferase as was previously accepted and highlights the versatility of the key enzymes in methane production in anoxic sediments.

## Introduction

Dimethylsulfide (DMS) is one of the most abundant volatile organic sulfur compounds with an estimated global production of over 300 million tons each year ^1^. DMS is also the largest source of biogenic sulfur in the atmosphere, where its oxidation products aid in cloud condensation and influence the atmospheric chemistry and potentially the Earth’s climate ^2^.

The main precursor of DMS in the environment is dimethylsulfoniopropionate (DMSP), an osmolyte produced in significant quantities (∼10^9^ tonnes annually) by marine algae, phytoplankton and plants such as *Spartina* and sugar cane ^3^. Recent studies have shown that bacteria are also important DMSP producers in both oxic and anoxic coastal and marine sediments ^4, 5^, suggesting these ecosystems to be highly productive environments for DMS production. Other key sources of DMS in sediments are the degradation of sulfur-containing amino acids and methoxylated aromatic compounds, reduction of dimethyl sulfoxide as well as the methylation of hydrogen sulfide and methanethiol (MT)^6, 7^.

In anoxic sediments, DMS can be degraded to potent greenhouse gases methane and carbon dioxide (CO_2_) by methylotrophic methanogens, further highlighting the significance of DMS^8. Cult^ivation-based studies on DMS-dependent methanogenesis showed that this process is carried out by certain methanogens of the genera *Methanomethylovorans*, *Methanolobus*, *Methanosarcina* and *Methanohalophilus* ^9–12^. In sulfate-containing environments, sulfate-reducing bacteria (SRB) of the genera *Desulfotomaculum* and *Desulfosarcina* can also use DMS as a carbon source ^13, 14^.

Despite the environmental significance of DMS and its degradation products (methane and CO_2_), the metabolic pathways of DMS-dependent methanogenesis have received little research interest. The accepted view is that, during methylotrophic methanogenesis, specific methyltransferases are used for each methylated compound (e.g. DMS, trimethylamine, methanol) although their substrate specificities have not been studied extensively ^15, 16^. There are only few studies on the metabolic pathways of DMS-dependent methanogenesis, which used pure cultures of *Methanosarcina barkeri* and *Methanosarcina acetivorans*, and suggest that methanogenesis from DMS is catalysed by methylthiol-CoM methyltransferase composed of two subunits (MtsA and MtsB) ^16, 17^. Also, MtsD, MtsF and MtsH were designated as methylsulfide-specific methyltransferases with central roles of MtsF and MtsH in *M. acetivorans* to produce methane from DMS ^18^. Conversely, Fu and Metcalf (2015) showed that MtsD is the specific methyltransferase in *M. acetivorans* to carry out DMS- dependent methanogenesis ^17^. Nevertheless, the metabolic pathways of DMS-dependent methanogenesis in the environment is undocumented.

Here, we studied the microbial diversity and metabolic pathways underpinning DMS-dependent methanogenesis in anoxic sediments from the Baltic Sea. Permanently hypoxic or anoxic conditions as well as the brackish nature of the Baltic Sea sediments provide an ideal ecosystem to study anaerobic DMS degradation ^19^. Our approach combining anoxic sediment incubations, amplicon sequencing, genome-centric metagenomics and metatranscriptomics provide the first insight into the sediment depth-profile of the methanogen diversity and key enzymes in DMS-dependent methanogenesis.

## Materials and Methods

### Study area and sampling

The study sites were located in Himmerfjärden, Baltic Sea, Sweden (Supplementary Figure 1). The bay has a salinity between 5 and 7‰, and consists of a series of small depositional basins with maximum water depths between 25 m and 50 m that accumulated fine-grained organic-rich sediment. Organic carbon concentrations in the investigated sediments vary between 3 and 4% dry weight ^20^. The depth of the sulfate-containing sediments vary between 25 cm and 40 cm depending on season ^21^. Below this depth, sediments show high rates of methanogenesis leading to the accumulation of methane ^20^. Bottom waters in the lowermost meter of the bay are oxic or hypoxic throughout the year with concentrations generally above 60 µmol L^-1^. However, oxygen uptake rate are high so that oxygen penetration depths are only between 0.24 cm and 0.63 cm ^22^.

Three sites were sampled using the research vessel R/V Limanda: Station H2 (N58°50’55, E17°47’42), H3 (N58°56’04, E17°43’81) and H5 (N59°02’21, E17°43’59). Duplicated sediment cores were collected using a multicorer (40 cm) and a small Rumohr-type gravity corer (140 cm). The sediment cores were transported to the Askö Laboratory of the Stockholm University Baltic Sea Centre and sliced into seven layers according to the sulfate concentrations of the sediment pore water (0 and 4.5 mM; Table 1) ^20^. The sediment slices were vacuum sealed into gas-tight bags and transported to Queen Mary University of London the next day in a cool box kept below 8°C. Incubations were set up the same day and a portion of each sediment layer was placed at -20°C for DNA extraction.

**Table 1.**
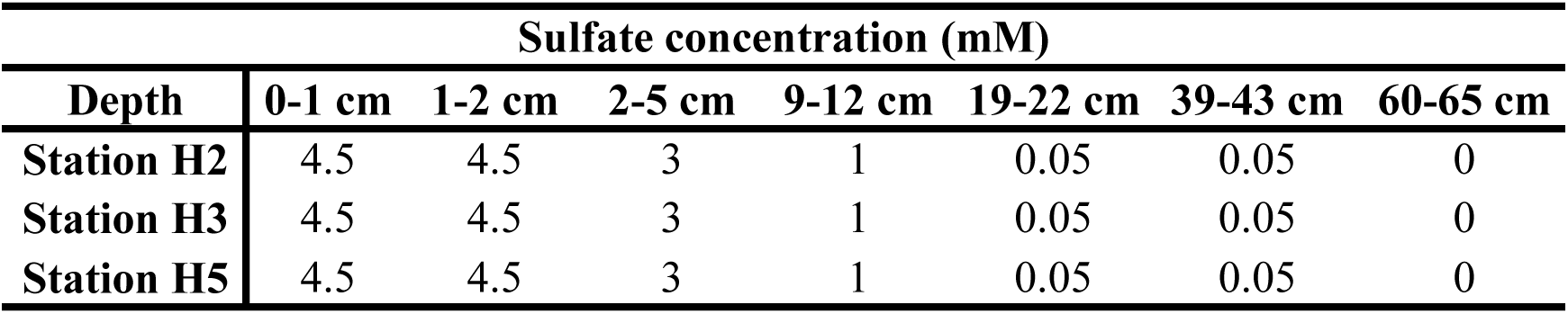
Sulfate concentrations in sediment layers from the three sampling stations. These also represent the sulfate concentrations provided in the incubations.

| Sulfate concentration (mM) |  |  |  |  |  |  |  |
| --- | --- | --- | --- | --- | --- | --- | --- |
| Depth | 0-1 cm | 1-2 cm | 2-5 cm | 9-12 cm | 19-22 cm | 39-43 cm | 60-65 cm |
| Station H2 | 4.5 | 4.5 | 3 | 1 | 0.05 | 0.05 | 0 |
| Station H3 | 4.5 | 4.5 | 3 | 1 | 0.05 | 0.05 | 0 |
| Station H5 | 4.5 | 4.5 | 3 | 1 | 0.05 | 0.05 | 0 |

### Incubation set-up

Triplicate incubations were prepared in an anaerobic glove box (Belle Technology, UK) for each sampling location and depth using 2.5 g of homogenised sediment and 20 mL of artificial seawater (ASW). Two sets of replicated controls were also set up. One set contained no DMS, and the other contained DMS and triple autoclaved sediment to monitor any sediment adsorption of DMS. The ASW consisted of 0.32 M NaCl, 10 mM MgSO_4_.7H_2_O, 8.8 mM NaNO_3_, 3.1 mM CaCl_2_.2H_2_O, 10 mM MgCl_2_.6H_2_O, 9 mM Trizma base, 0.21 mM K_2_HPO_4_.3H_2_O, trace elements, and vitamins ^23^. The sulfate concentrations of the incubations were adjusted according to the *in situ* sulfate concentration of the sediment pore water at each depth (Table 1). The microcosms were incubated in the dark and at 8°C to avoid the photochemical destruction of DMS ^24^.

Each sample was amended with DMS as the carbon and energy source. Initially, samples were amended with 2 µmol g^-1^ DMS. After the initial DMS degradation, another 2 µmol g^-1^ DMS were added. All subsequent additions were 4 µmol g^-1^ DMS. The incubations were terminated when cumulative methane concentrations became stable (between 82 and 128 days), which corresponded to the total DMS additions of 9.7 – 51.9 µmol g^-1^ DMS. After the incubation period, the supernatant and sediment were separated via centrifugation at 1,000 rpm for 6 min, and placed at -20°C and -80°C, respectively, until further analysis.

### Analytical measurements

DMS in the incubation bottles was measured on a gas chromatograph (GC; Agilent Technologies, 6890A Series, USA) fitted with a flame photometric detector (FPD) and a J&W DB-1 column (30 m x 0.32 mm Ø; Agilent Technologies, USA). The oven temperature was 180°C, and zero grade N_2_ (BOC, UK) was the carrier gas (26.7 mL min^-1^). FPD was run at 250°C with H_2_ and air (BOC, UK) at a flow rate of 40 and 60 mL min^-1^, respectively. DMS standards (50 µM – 10 mM) were prepared by diluting >99% DMS (Sigma-Aldrich, USA) in distilled anoxic water previously prepared by flushing with oxygen-free N_2_ (BOC, UK).

Methane and CO_2_ were measured using GC (Agilent Technologies, USA, 6890N Series) fitted with a flame ionisation detector (FID), Porapak (Q 80/100) packed stainless steel column (1.83 m x 3.18 mm Ø; Supelco, USA), and hot-nickel catalyst which reduced CO_2_ to methane (Agilent Technologies, USA). The oven temperature was 30°C, and zero grade N_2_ (BOC, UK) was the carrier gas (14 mL min^-1^). FID was run at 300°C with H_2_ and air (BOC, UK) at a flow rate of 40 and 430 mL min^-1^, respectively. The GC was calibrated against certified gas mixture standards (100 ppm methane, 3700 ppm CO_2_, 100 ppm N_2_O, balance N_2_; BOC, UK). The total methane concentrations in the incubation bottles also included dissolved methane in ASW calculated using the atmospheric equilibrium solubility equation as a function of temperature, salinity and headspace concentration ^25^.

The total CO_2_ production was the sum of the CO_2_ concentration in the headspace and the total dissolved inorganic carbon (DIC) in the water phase. The CO_2_ in the headspace was measured using a gas chromatograph as above. Total DIC was measured as CO_2_ in the headspace after the supernatant of the slurry was fixed with 24 µL ZnCl_2_ (50% w/v) and acidified with 100 µL 35% HCl. An inorganic calibration series (0.1-8 mM) of Na_2_CO_3_ was used as a standard (Sigma-Aldrich, USA).

Sulfate concentrations were measured at the end of incubation, using porewater filtered through 0.2 μm syringe filters (PTFE hydrophilic; Fisher Scientific, USA). An ICS-5000 Dual Gradient RFIC Ion Chromatograph (Thermo Fisher Scientific, USA) equipped with a Dionex IonPac AS11-HC-4 μm column (2x250 mm) and a Dionex IonPac AG11-HC-4 μm guard column (2x50 mm) was used. A gradient of 1.5-22 mM KOH (Dionex EGC 500 KOH, with CR-ATC column) was applied as eluent.

### DNA extraction, PCR and quantitative PCR

DNA was extracted from both the original and DMS-incubated sediment samples using the DNeasy Powersoil kit (Qiagen, NL) following the manufacturer’s instructions using FastPrep96 (MP Biomedicals, USA) at maximum speed for 40 sec. The *mcrA* gene, which encodes the *α*-subunit of the methylcoenzyme M reductase in all known methanogens, was amplified using the mcrIRD primer set ^26^. The PCR solution contained 1 µL of DNA template, 1 µL of each primer (10 µM), 25 µL 2x MyTaq HS Red Mix (Meridian Bioscience, USA), and 22 µL ultra-pure water. PCR conditions were 95 °C for 5 min and 39 cycles of 95 ^oC for^ 1 min, 51 °C for 1 min, 72 °C for 1 min and a final extension at 72 °C for 5 min. PCR products were cleaned using JetSeq Clean beads (1.4x; Meridian Bioscience, USA) following the manufacturer’s instructions.

Quantitative PCR (qPCR) of the *mcrA* gene was carried out in triplicate using the primers mlas-mod-F and mcrA-rev-R, a CFX384 Touch Real-Time PCR Detection System (Bio-Rad Laboratories, USA) and a low volume liquid handling robot for automation (Mosquito HV, SPT Labtech, UK) ^27, 28^. Each reaction contained 4 ng/µL DNA, 10 µM of each primer, 2.5 µL SensiFAST SYBR (No-ROX; Meridian Bioscience, USA), and 1.8 µL ultra-pure water. The cycling conditions were 95°C for 3 min, followed by 40 cycles of 95°C for 15 sec, 65°C for 30 sec, and 72°C for 20 sec. A melt curve analysis was performed by increasing the temperature from 65°C to 95°C in 0.5°C increments. Standard curves were produced using a serial 10-fold dilution of clones containing the *mcrA* gene. The reaction efficiency was between 90% and 110%, and the R^2^ value for the standard curve was >99%.

### High-throughput sequencing and sequence analysis

For the sequencing library preparation, a second PCR was carried out to attach overhang adapters to the cleaned-up PCR products using *mcrA* primers containing 5’ overhang adapters (10 µM). The PCR conditions were 95°C for 3 min, 15 cycles of 95°C for 20 sec, 55°C for 15 sec, 72°C for 15 sec and a final extension step at 72 °C for 5 min. After clean-up, the PCR products were further amplified for the addition of dual indices using 2 µL of the clean barcoded PCR products, 1 µL of each primer (5 µM), 12.5 µL 2x Q5 Hot-start Ready mix (NEB, USA), and 8.5 µL ultra-pure water. The PCR conditions were as above. All PCR products were normalized using the SequalPrep Normalization Plate kit (Invitrogen, USA) and sequenced on a MiSeq Next Generation sequencing platform (2x300 bp; Illumina; USA).

The amplicon sequences were analysed using QIIME2 2021.11 on Queen Mary University of London’s Apocrita HPC facility, supported by QMUL Research-IT ^29, 30^. Taxonomy was assigned to Amplicon Sequence Variants using Naive Bayes classifiers, trained using a custom *mcrA* database compiled using FunGenes, Python 3.10.8 and the RESCRIPt package in QIIME2 ^31, 32^.

### Statistical analysis

All statistical analyses including the calculations of the Shannon index, permutation tests of multivariate homogeneity of group dispersions (999 permutations), ANOVA, pairwise PERMANOVA (9999 permutations) and the principal coordinate analyses (PCoA) with Bray-Curtis dissimilarity were carried out and visualised using microeco and ggplot2 in RStudio (2022.07.1) ^33–35^. Spearman’s correlation analysis (r_s_) between the first three PCoA coordinates and the consumed DMS, produced methane and CO_2_, and sulfate concentrations was conducted using PAST 4.2 ^36^.

### Metagenomics analysis

Paired-end (2x150 bp) metagenomics sequencing of the DMS-incubated sediments from 19- 22 cm depth from the three stations was conducted on the Illumina NovaSeq 6000 platform at U.S. Department of Energy (DOE) Joint Genome Institute (JGI). The quality of the DNA was analysed using Nanodrop One (Thermo Scientific, USA) and Qubit 2.0 Fluorometer (Invitrogen, USA). The A_260/280_ ratio of the samples was between 1.6 and 2.0, whereas the A_260/230_ ratio was between 1.8 and 2.2. The DNA concentrations were between 10-15 ng/µL. In total, 154 Gb of sequencing data corresponding to 64.5 Gb from station H2, 41.7 Gb from H3 and 47.9 Gb from H5, were obtained. The data analysis was performed by JGI following the well-established JGI workflow, including assembly, feature prediction, annotation and binning ^37^.

Metagenome-assembled genomes (MAGs) were recovered using MetaBAT 2.12.1 ^38^. Genome completion and contamination were estimated using CheckM 1.0.12 ^39^. The genome bins were assigned as high (HQ) or medium quality (MQ) according to the Minimum Information about a Metagenome-Assembled Genome (MIMAG) standards ^40^. Integrated Microbial Genome (IMG) and GTDB-tk (0.2.2) databases were used to infer taxonomic affiliations ^41, 42^.

A list of 78 genes involved in methanogenesis (15 genes specific to methylotrophic methanogenesis) was compiled using the MetaCyc and KEGG databases (Supplementary Table 2) and quantified in the metagenomics datasets and metagenome assembled genomes (MAGs) ^43, 44^. All absolute abundance counts were normalised using the CPM (copies per million) normalisation method and log-transformed. R (4.2.1) and ggplot2 were used to make a heatmap showing the log_10(CPM)_ values for each gene ^33, 35, 45^.

### Metatranscriptomics analysis

Metatranscriptomics were conducted on the DMS-incubated sediments (19-22 cm depth) from the three stations. Total RNA was extracted using the ZymoBIOMICS RNA miniprep Kit (Zymo Research, USA) according to the manufacturer’s instructions. The quality was checked using a Tapestation 2200 (Agilent Technologies, USA) and the absorbance was measured using a Nanodrop One (Thermo Scientific, USA). The A_260/280_ ratio was greater than 1.8 in all samples. The concentration of total RNA ranged between 17 ng/µL and 88 ng/µL (Qubit 2.0; Invitrogen, CA, USA). DOE JGI performed the metatranscriptomics sequencing (2x150 bp) on the Illumina NovaSeq S4 platform and analysed the sequences following a well-established JGI-created workflow. Metatranscriptomics sequencing of the sample from station H2 was not successful. In total, 98.1 Gb of sequencing data corresponding to 44.4 Gb for the H3 and 53.7 Gb for the H5 sample, were obtained.

A total of 78 methanogenesis-related genes (Supplementary Table 2) were screened within the metatranscriptomes. Pyrrolysine, an in-frame amber codon (UAG), which does not act as a stop codon during synthesis, was searched within the methyltransferase gene fragments using JGI’s Chromosome Viewer ^46^. If one fragment contained pyrrolysine, the two fragments were merged. Then, the absolute abundance of the genes was calculated, normalised and log-transformed. Fragments per kilobase of transcript per million fragments mapped reads (FPKM) was calculated and a heatmap was created using ggplot2 ^35, 47^.

### *Methanolobus* genome analysis for the *mts* genes

A total of 17 complete *Methanolobus* genomes were collected from NCBI and JGI IMG/MER databases ^41, 48^. Using protein Basic Local Alignment Search Tool, these genomes were screened for the presence of the Mts proteins (Q48924, Q8PUA8, Q8PUA7, AAM04298.1, WP_048180685.1, AAM07726, WP_048180700.1, AAM07897.1, WP_048177073.1) downloaded from the Uniprot and NCBI databases ^48, 49^.

### Phylogenetic analysis

The PhyloFunDB pipeline was used for the construction of the phylogenetic tree of the *mcrA* gene. The *mcrA* sequences from uncultured methanogens were manually removed and those from *Methanolobus oregonensis*, *Methanolobus taylorii* and *Methanolobus tindarius* were added. IQ-TREE (1.6.12) was used to create the phylogenetic tree using *Methanopyrales* as the out-group with 1000 bootstrap replications ^50^. ModelFinder was used to find the best-fit model (mtZOA+F+G4) ^51^.

Phylogenetic analyses of methyltransferase and corrinoid proteins were performed in MEGA7 using the neighbour-joining method with 100 bootstrap replications and the Poisson correction method ^52^.

## Results

### Sediment depth profiles of DMS, methane and CO_2_

We terminated the incubations between day 82 and 128, when cumulative methane concentrations became stable (Supplementary Figure 2). We observed a lag phase in all incubations before methane production was detected although the DMS degradation began within the first couple of days, suggesting that SRB started to consume DMS before methanogens. We observed DMS degradation and accompanying methane and CO_2_ productions in all sediment incubations except for the 60-65 cm (bottom) sediment layer from the H2 station, where no methane production was observed despite DMS degradation.

The greatest DMS consumption and methane production were recorded in the top two sediment layers (0-1 cm and 1-2 cm; Figure 1) from the H3 station although these incubations had the highest initial sulfate concentration (5 mM; Supplementary Figure 3). The DMS consumption was 39.1±4 and 48.5±2 μmol g^-1^ wet sediment, whilst the net methane production was 44.9±11.6 and 62.9±1.8 μmol methane g^-1^ wet sediment, respectively, which correspond to 77% to 86% of the theoretical methane yield (58.6±6 and 72.7±3 μmol g^-1^ wet sediment) assuming 1 mol of DMS yields 1.5 mol of methane ^53^. This indicates ∼30 and ∼41 μmol DMS g^-1^ wet sediment were converted to methane in these samples, suggesting the rest of the DMS (∼9 and ∼7.5 μmol DMS g^-1^, respectively) was degraded by SRB. Supporting this, ∼77% of the sulfate amended to these incubations was consumed, decreasing the sulfate concentration to 9±0.2 μmol g^-1^ wet sediment (Supplementary Figure 3).

**Figure 1.**
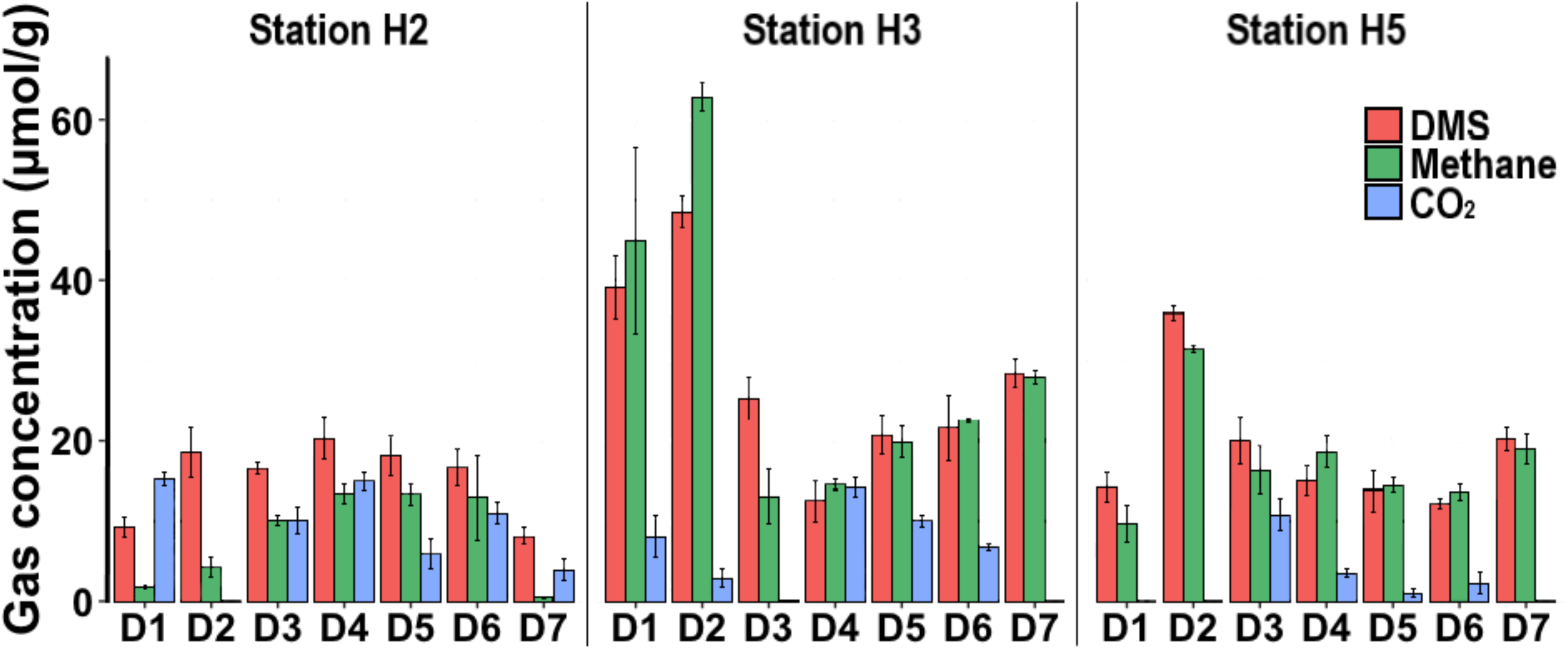
Average of total amount of degraded DMS, methane and CO_2_ per gram of DMS- amended sediment after 82-128 days of incubation. D1: 0-1 cm; D2: 1-2 cm; D3: 2-5 cm; D4: 9-12 cm; D5: 19-22 cm; D6: 39-43 cm; D7: 60-65 cm.

DMS degradation and net methane production were comparatively low in H2 sediment incubations, where the maximum methane production (13±2.6 μmol g^-1^) was observed in sediments from 9-12 cm, 19-22 cm and 39-43 cm of depth. Stoichiometrically, this corresponds to ∼8.5 μmol DMS g^-1^ consumption, however, the actual concentrations of degraded DMS were 20.3±2.5, 18.1±2.5 and 16.6±2.3 μmol DMS g^-1^, respectively (Figure 1). Similarly, the highest methane production was 31.4±0.4 μmol g^-1^ in the H5 sediment incubations of the 1-2 cm depth interval. This methane production corresponds to ∼21 μmol DMS g^-1^ degradation, however, a total of 35.9±0.9 μmol DMS g^-1^ was degraded in these incubations. These results indicate that part of the DMS was degraded via the sulfate reduction route in these incubations. Intriguingly, the sulfate concentrations in the sediments below 9 cm from all three sites increased significantly compared to the initial concentrations (Supplementary Figure 3). This suggests that hydrogen sulfide produced as one of the end products of DMS degradation was converted to sulfate, which led to cryptic sulfur cycling in these incubations ^8^.

We also measured CO_2_ in the incubations as it is one of the metabolic end products of anaerobic DMS degradation via both methanogenesis and sulfate reduction (Figure 1). In general, the total amount of CO_2_ was significantly lower than the theoretical CO_2_ amounts assuming only methanogenesis or sulfate reduction took place (2 mol and 0.5 mol per mol of DMS, respectively), implying that CO_2_ was simultaneously consumed in our incubations ^13, 53^.

### Depth profiles of methanogen diversity and abundance

We characterised the depth profiles of methanogen diversity and abundance in our original and DMS-amended sediment samples via sequencing and quantifying the *mcrA* gene.

There was a statistically significant difference in methanogen diversity between the original and DMS-amended sediment samples (PERMANOVA; p<0.01), whilst there was no difference between the control incubations without DMS and the original sediments, indicating that the shift in methanogen diversity was due to DMS addition.

All original Baltic Sea sediment samples from the surface down to the bottom of the sulfate-methane transition zone (SMTZ) at19 cm, had strong dominance of *Methanolobus* (47%- 80%; Figure 2a). Below this depth, the methanogen diversity in the original sediments becomes more varied with *Methanoculleus* (37%-75%), unclassified Archaea (1%-36%) and Candidatus *Methanomethylophilus* (3%-42%) in addition to *Methanolobus* (3%-37%), highlighting a shift in methanogen populations below the SMTZ.

**Figure 2.**
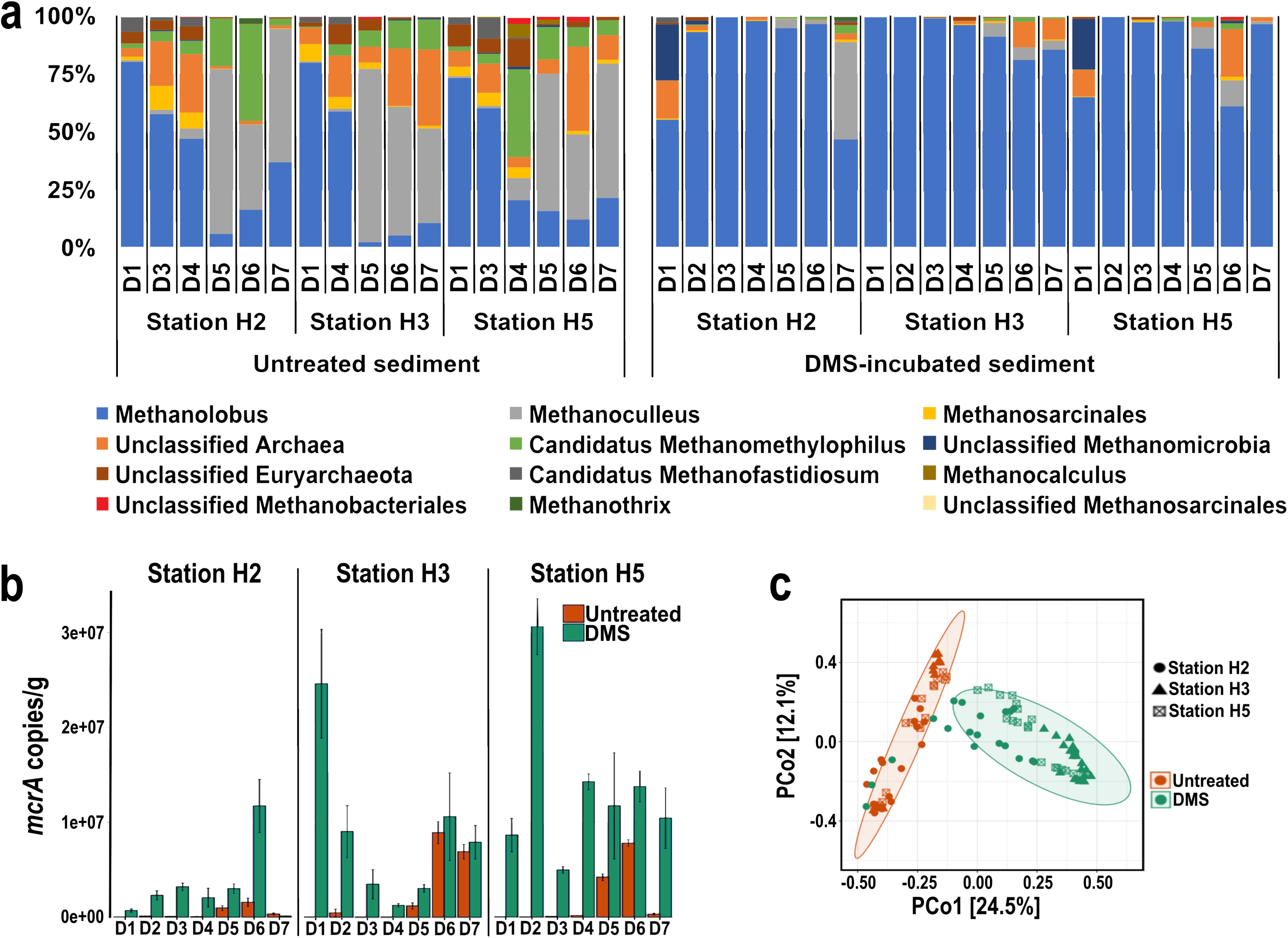
a) Relative abundance of methanogens at genus level based on *mcrA* sequencing. *Methanolobus* dominated in all original sediment down to 19 cm (D1-D4) and DMS- amended sediment except for sample H2D7 where methane production was minimal. b) the *mcrA* gene copy numbers in original and DMS-amended sediment incubations as determined by qPCR. All reactions were set up in triplicate and the average abundance and standard error were shown. c) PCoA plot of the *mcrA* sequences based on Bray-Curtis dissimilarity metrics. Ellipses indicate 95% confidence intervals according to treatment data. Colour indicates treatment (red: untreated; green: DMS-amended). Shapes indicate sampling site. D1: 0-1 cm; D2: 1-2 cm; D3: 2-5 cm; D4: 9-12 cm; D5: 19-22 cm; D6: 39-43 cm; D7: 60-65 cm.

All DMS-amended samples, except for the H2 and H5 top and H2 bottom layers, where low or no methane production was observed, had a sharp increase in the relative abundance of *Methanolobus* (61 – 99%) regardless of the sulfate concentration in the incubations (Figure 2a). In the H2 and H5 top sediment incubations, unclassified *Methanomicrobia* increased to 24%±3% and 22%±5%, respectively, although this taxon was not detected in the original sediment samples. To assess the factors influencing the methanogen diversity in the original and DMS-amended sediments, we conducted a principal coordinate analysis, which clearly separated the original and DMS-amended sediment samples (Figure 2b). Spearman’s correlation analysis of the first principal coordinate (explaining 24.5% of the total variation in methanogen community composition) correlated positively with DMS degradation, methane and CO_2_ production (Supplementary Table 1; p<0.001).

The abundance of methanogens increased significantly in DMS-amended incubations, where methane production was observed compared to the original sediment samples (Supplementary Figure 4, p<0.05). However, the correlation between the *mcrA* abundance and the methane production was not linear.

### Taxonomic analysis of metagenomes from DMS-amended incubations

To gain further insight into the microbial populations degrading DMS, we conducted metagenomic sequencing of the DMS-incubated sediments from all three stations at the SMTZ (19-22 cm), where both DMS-dependent methane production and sulfate reduction are likely to happen *in situ*.

Taxonomic classification of the metagenomes showed that *Methanolobus* were the dominant methanogen (69%-87% amongst Archaea) in all three DMS-incubated samples. We also analysed the *mcrA* sequences retrieved from the assembled metagenomes, which indicated that 35% of the sequences were most closely affiliated with *Methanolobus* (89.6% to 98.8% similarity; Figure 3). Furthermore, we successfully constructed 44 MAGs from the metagenomes (Supplementary Table 3). Four of these MAGs were methanogens recovered from the three stations. These medium quality MAGs have completeness ranging between 62.5%-92.8% and contamination <2%, but they do not contain all three rRNA genes ^40^. We retrieved one *mcrA* sequence from the methanogen MAG (92% completeness) obtained from the H2 sample (19-22 cm) and it was most similar to the *mcrA* sequence from *Methanolobus vulcani* (WP_091708234; 94.6% amino acid similarity). The phylogenetic analysis shows all *mcrA* sequences in the metagenome datasets and the MAG clustered together with *mcrA* sequences from cultured *Methanolobus* species (Figure 3).

**Figure 3.**
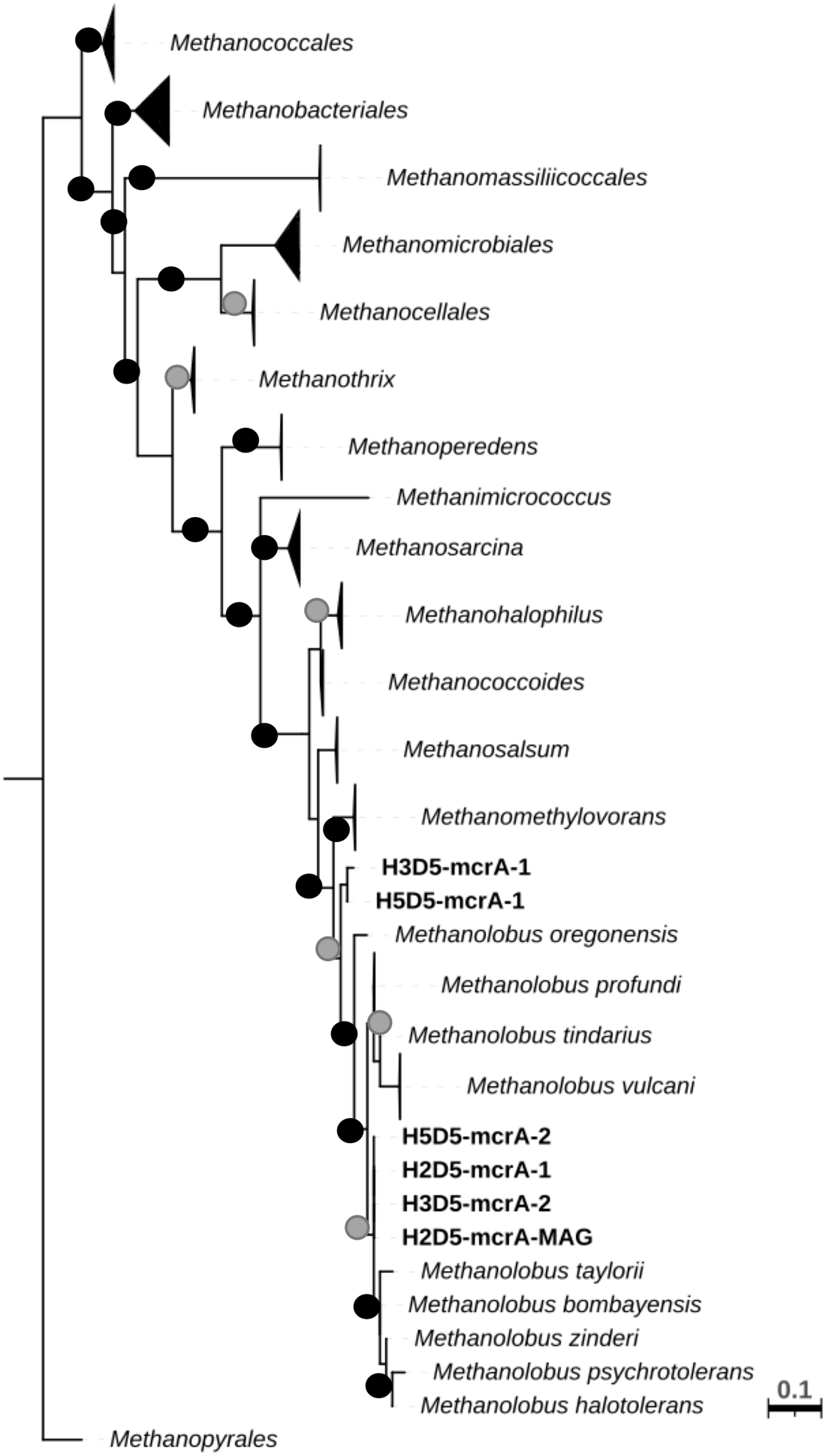
Maximum likelihood phylogenetic tree of the *mcrA* gene from the cultured methanogens. The tree also contains the *mcrA* sequences (marked in bold) from the metagenomes and the MAGs obtained within this study. ModelFinder was used to find the best-fit model for the data ^51^. Bootstrap values (1000 replicates) were shown as black dots (>80%) and grey dots (<80%). The tree is drawn to scale, with branch lengths accounting for substitutions per site. The genus *Methanopyrales* was used as the outgroup.

### Metabolic pathways of DMS degradation in the sediment incubations

We analysed the metabolic pathways of anaerobic DMS degradation via metagenome and metatranscriptome analyses of the samples from DMS-amended incubations with sediments at 19-22 cm. We screened for 78 genes involved in methane production in the metagenomics and metatranscriptomics datasets, and also the constructed methanogen MAGs (Supplementary Table 2 and 3).

Notably, the relative expressions of the *mtsA, mtsB and mtsH* genes encoding for MT- and DMS-methyltransferases characterised in *M. barkeri* and *M. acetivorans* were low (<0.01%) or even undetectable in the metatranscriptomics datasets (Figure 4a). *mtsD* and *mtsF* had higher abundances than the other *mts* genes (0.3% and 0.1%, respectively), however they were identified as *Methanosarcina, Methanomethylovorans* and *Methanococcoides* with similarities between 82%-98%. Similarly, these genes were absent or in low abundance (<0.03%) in the metagenomes, and were affiliated with *Methanomethylovorans* (Figure 4b). Furthermore, we did not find these genes in the four methanogen MAGs (Supplementary Figure 5). Surprisingly, the relative expression of the genes encoding for trimethylamine (TMA)- and methanol-corrinoid protein co-methyltransferases (*mttB* and *mtaB*, respectively) were dramatically high at 5.4% and 9% relative expressions. Moreover, the relative expressions of the whole gene clusters encoding dimethylamine TMA- and methanol- methyltransferases (*mttBC* and *mtaABC*, respectively) were higher (3-3.5% and 3.5-4.2%, respectively) than that of the *mts* gene cluster (<0.002%; Figure 4). The genes encoding for TMA- and methanol-methyltransferases were also present in the metagenomes from all three samples. *mttB*, encoding for TMA-corrinoid protein co-methyltransferase, was the most abundantly found gene (∼6%) involved in methylotrophic methanogenesis (Figure 4b). This was significantly higher than all other methylotrophic methanogenesis genes searched (p<0.001; Figure 4b).

**Figure 4.**
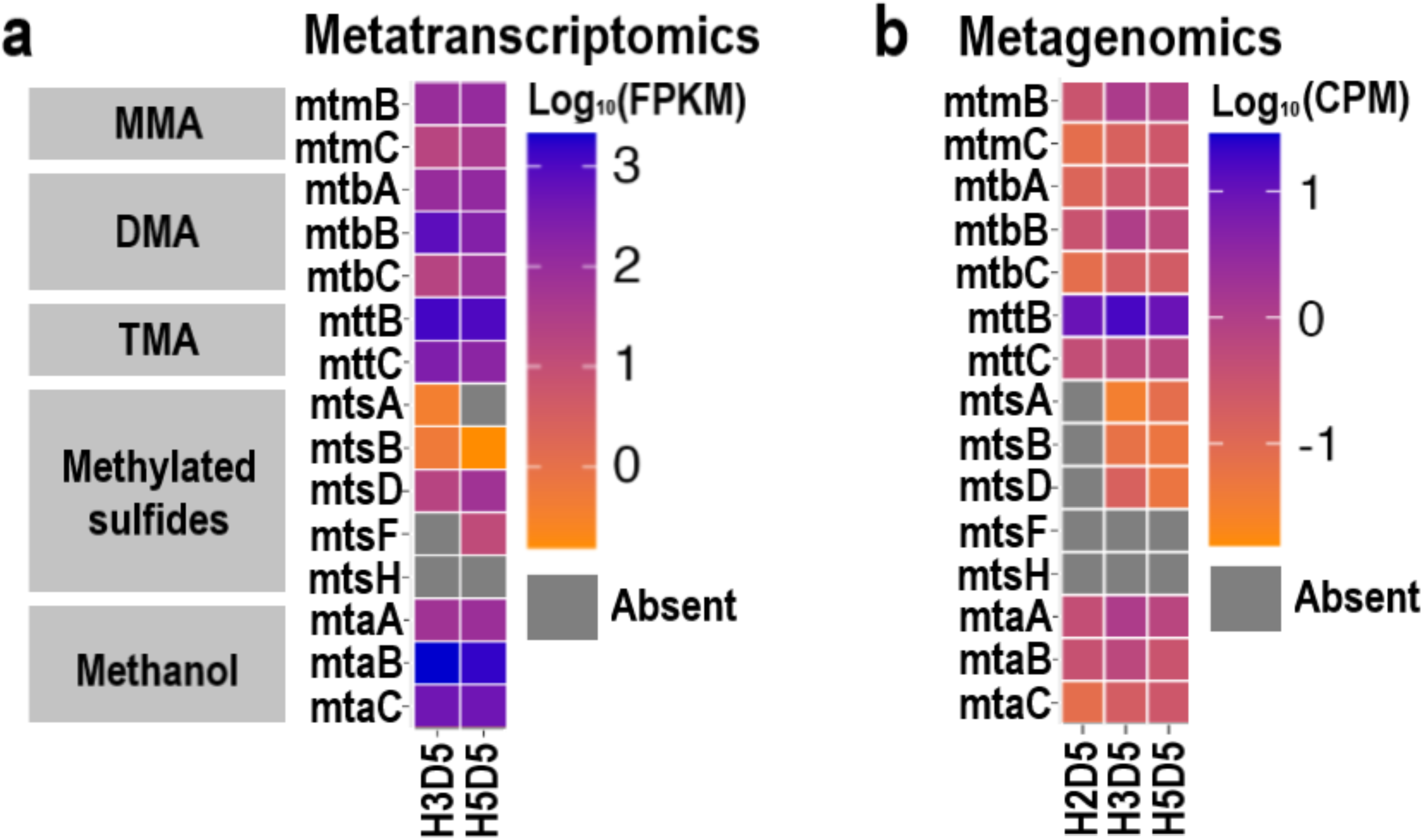
Heatmaps showing expression and abundance of genes involved in methylotrophic methane production. a) Metatranscriptomics datasets; b) Metagenomics datasets. FPKM: fragments per kilobase of gene per million reads, CPM: Copies per million reads.

The taxonomic profiling of the genes encoding for TMA and methanol methyltransferases and corrinoid proteins (MtaB, MtaC, MttB, MttC) from the metatranscriptome datasets assigned them to *Methanolobus* (Figure 5). In line with this and the metagenomics sequence analysis, the entire gene clusters encoding for DMA-, TMA- and methanol- methyltransferases (*mtbABC*, *mttABC* and *mtaABC*, respectively) were also present in the two most complete *Methanolobus* MAGs (H2D5-*Methanolobus* and H5D5-*Methanolobus*; Supplementary Figure 5).

**Figure 5.**
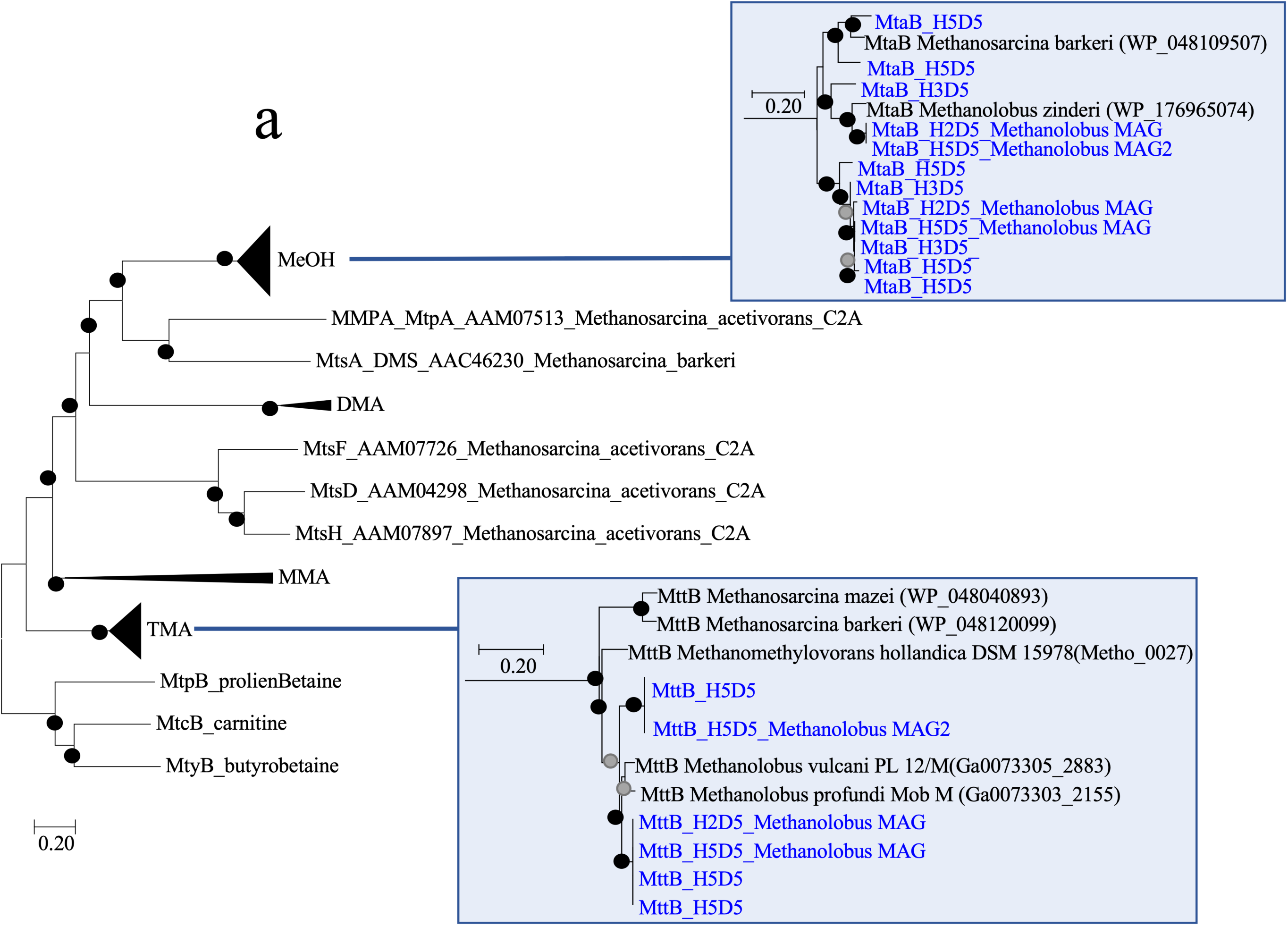
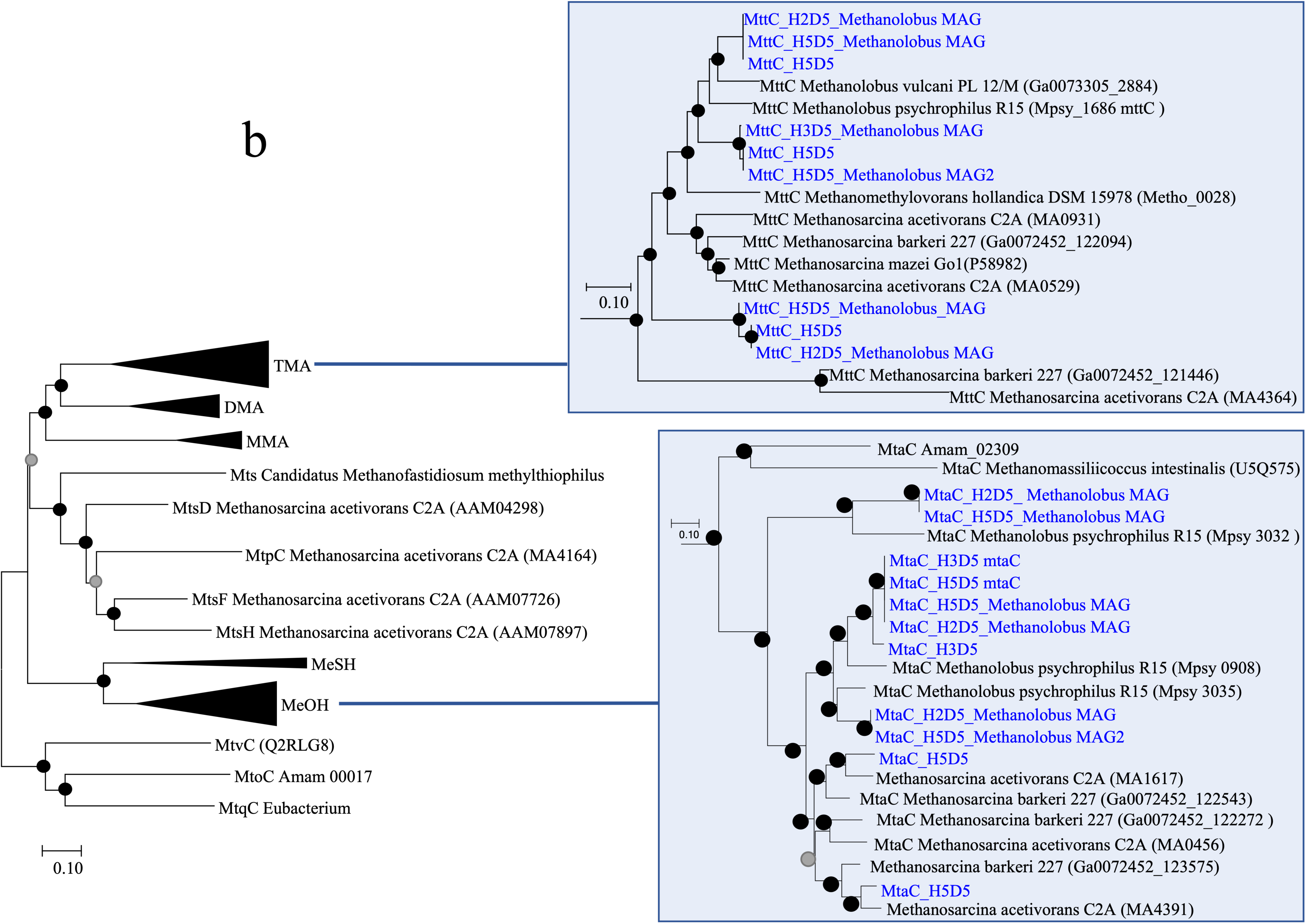
Phylogenetic tree of (a) MT1 methyltransferase and (b) corrinoid proteins including sequences from metatranscriptomics and MAGs recovered from the metagenomics datasets. The evolutionary history was inferred using the Neighbour-Joining method. The optimal trees with the sum of branch length of 11.7 and 11.1 are shown for methyltransferase and corrinoid proteins, respectively. Bootstrap values (100 replicates) are shown as black (>50) and grey (<50%) dots. The tree is drawn to scale, with branch lengths in the same units as those of the evolutionary distances used to infer the phylogenetic tree. All ambiguous positions were removed for each sequence pair. There was a total of 637 positions for methyltransferases and 217 positions for the corrinoid proteins in the final dataset.

We also searched for genes encoding for key enzymes common to all methanogenic pathways (Supplementary Table 2). We found that all the genes in the *mcrABCDG* operon had a relative expression of >1% in metatranscriptomics datasets (Supplementary Figure 6a). We further showed that the transcripts of several other gene clusters in central methanogenic pathway (e.g. *mtrA-H*, *hdrA-D*, *mvdADG*, *frhABDG*; Supplementary Table 2) were found at levels 0.26%, 1.75% and 0.84%, respectively. *hdrA* was found at strikingly high level (6.65%) compared to others, which is likely because this gene is conserved across all methanogens ^54^. On the other hand, *fpo* and *vho* genes catalysing coenzyme B/coenzyme M regeneration were not transcribed in our sediment incubations. These genes were also absent in the metagenomics datasets (Supplementary Figure 6b).

To understand whether acetoclastic and hydrogenotrophic methanogenesis pathways were active in our DMS incubations, we searched for genes specific to these methanogenesis pathways (*ack*, *acs*, *coo*, *cdh*, *pta* for acetoclastic and *fmd*, *ftr*, *mch*, *mer* for hydrogenotrophic methanogenesis). All genes except for *acs* were expressed at <0.1%, while *acs* was expressed at 4.9% (Supplementary Figure 7a). It should, however, be kept in mind that methylotrophic methanogens also possess the *acs* gene ^55^.

## Discussion

Despite the environmental importance of DMS as a methane precursor in anoxic sediments, limited information concerning the microbial diversity and metabolism of DMS-dependent methanogenesis is available. Here, we conducted the first study on the depth profile of the microbial populations and metabolic pathways underlying DMS-dependent methanogenesis in anoxic sediments.

Our sediment incubations have shown that DMS degradation proceeds via both methanogenesis and sulfate-reduction throughout the sediment sampled at the three stations in the Baltic Sea. Higher methane yields from DMS degradation were observed in H3 and H5 stations. This may be due to higher inputs of organic carbon and nutrient from the discharge of an upstream sewage treatment plant, leading to higher rates of carbon mineralisation allowing ultimately methanogenesis to occur^20^.

Multiple lines of evidence obtained from the amplicon sequencing, genome-centric metagenomics and metatranscriptomics data pointed that *Methanolobus* were the dominant DMS-degrading methanogens in our sediment incubations despite varying sulfate concentrations. This methanogen genus was also dominant in the original sediment samples, which suggests that halotolerant *Methanolobus* carry out methylotrophic methanogenesis in sulfate bearing sediments of the Baltic Sea and potentially degrade DMS when it is available. *Methanolobus* are known DMS degraders with several strains isolated from an oil well, marine, lake and estuarine sediments ^10, 11^. We also recently showed *Methanolobus* to be the dominant DMS-degrading methanogen genus in brackish sediments from the Medway Estuary, UK ^8^. Furthermore, a psychrotolerant *Methanolobus* strain has been isolated from a saline lake sediment in Siberia, indicating that this genus has members that can grow in low temperatures, as were measured in Baltic Sea sediments ^56^.

An important result of this study was the lack or very low detection (<0.3%) of the genes encoding for methylsulfide-methyltransferases (*mts*) in both metagenomics and metatranscriptomics sequences retrieved from incubations, where DMS-dependent methane production was observed. On the contrary, the transcriptional profiles of genes encoding for enzymes related to TMA- and methanol-methyltransferases (*mttB* and *mtaB*) showed much higher levels of gene transcription (5.4% and 9%, respectively) in DMS-amended incubations. These highly expressed methyltransferase genes were taxonomically affiliated with *Methanolobus*, supporting our findings via *mcrA* sequencing and taxonomic analysis of the metagenomes. We searched for the *mts* genes in all publicly available *Methanolobus* genomes and found that they do not contain the *mts* genes, further demonstrating that the *Methanolobus* genus do not use the *mts* genes when degrading DMS to methane.

Our results contradict previous studies, which proposed that each methylotrophic substrate requires a specific enzyme to methylate a corrinoid protein ^16, 17^. Furthermore, Tallant et al (2001) showed that the methylamine-specific methylcobalamin:CoM methlytransferase, MtbA, did not catalyse the methylation of cobalamine with DMS in *Methanosarcina barkeri*^57. Howe^ver, a recent survey analysing the presence of genes involved in methylotrophic methanogenesis within 465 metagenomes from wetlands, ocean and hypersaline sediments, showed a significantly low abundance of the *mtsA* compared to the *mttC* and *mtaA* that encode for TMA- and methanol-dependent methanogenesis genes, respectively ^58^. Given the high concentrations of DMS and its ubiquitous precursor DMSP in the environment, it is intriguing to count low levels of *mtsA* in environmental metagenomes. Hence, we propose that the *mtt* and *mta* genes, encoding for TMA- and methanol-methyltransferases, are versatile methyltransferases that can catalyse the transfer of the methyl moity of DMS to a corrinoid protein. This, however, does not exclude the possibility that there are novel methylsulfide-specific methyltransferases yet to be discovered.

In conclusion, our results provide the first evidence that DMS can be anaerobically degraded to methane via the activity of TMA and methanol methyltransferases in *Methanolobus*. This finding challenges the view that substrate-specific methyltransferases are used in methylotrophic methanogenesis. In light of the significance of this methanogenesis route in coastal and marine ecosystems, it is vital that the metabolic pathways underlying methylotrophic methanogenesis and the regulation of these pathways are unearthed.

## Data availability

All sequence data produced in this study are publicly available. The *mcrA* gene sequences are deposited at the National Center for Biotechnology Information (NCBI) Read Archive (PRJNA962783). Metagenomics and metatranscriptomics datasets are available at JGI GOLD database (Project IDs: Gp0507771, Gp0507772, Gp0507773, Gp0507777 and Gp0507778).

## Acknowledgements

This study was financially supported by UK Natural Environment Research Council (NE/S007725/1) and Queen Mary University of London with a postgraduate scholarship to S.L.T. The work (proposal:10.46936/10.25585/60001216) conducted by the U.S. Department of Energy Joint Genome Institute (https://ror.org/04xm1d337), a DOE Office of Science User Facility, is supported by the Office of Science of the U.S. Department of Energy operated under Contract No. DE-AC02-05CH11231.

