## Supplementray Material for "*Methanolobus* use unspecific methyltransferases to produce methane from dimethylsulfide"

### Supplementary Material

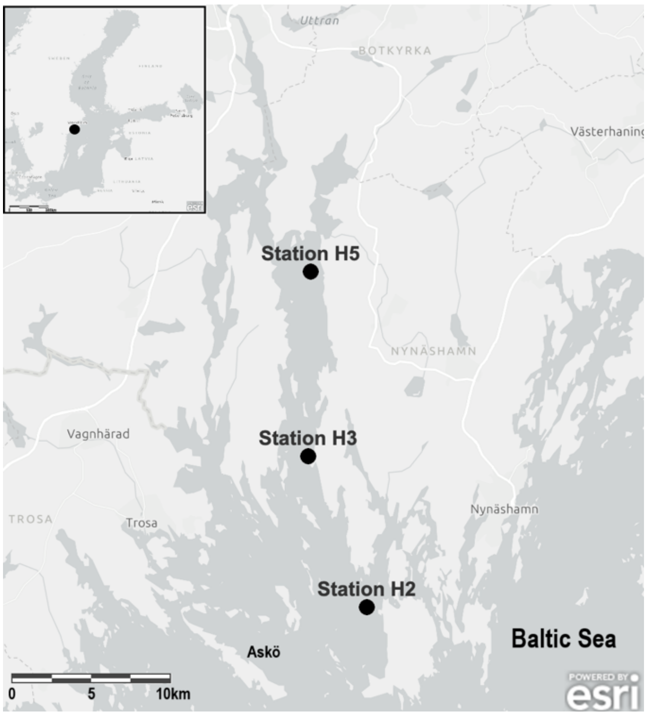


Supplementary Figure 1. Map of Himmerfjärden and the Baltic Sea showing the three sampling stations H2, H3 and H5. The Stockholm University Baltic Sea Centre is located on the Askö Island. Inset map shows the entire Baltic Sea.

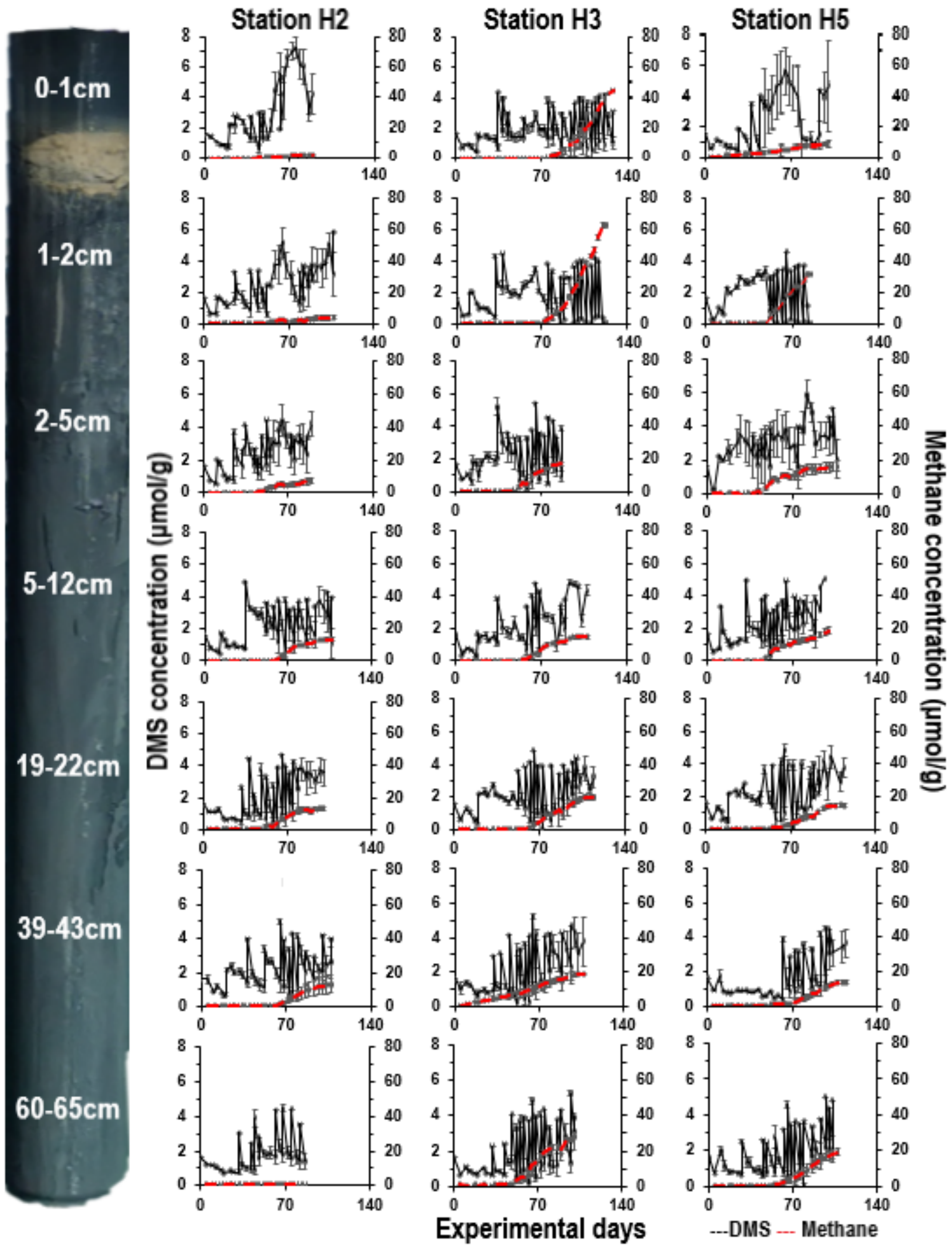


Supplementary Figure 2. Average concentrations of DMS and methane in DMS-amended incubations from seven sediment layers (0-1 cm, 1-2 cm, 2-5 cm, 5-12 cm, 19-22 cm, 39-43 cm, 60-65 cm). Black lines: DMS; Red lines: Methane

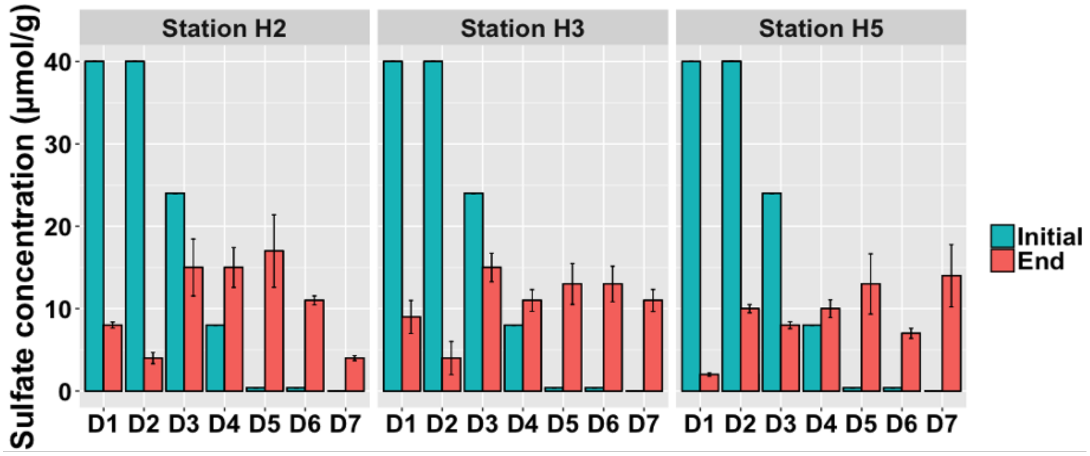


Supplementary Figure 3. Average sulfate concentrations at the start and the end of the incubation period in the samples D1: 0-1 cm; D2: 1-2 cm; D3: 2-5 cm; D4: 9-12 cm; D5: 19-22 cm; D6: 39-43 cm; D7: 60-65 cm.

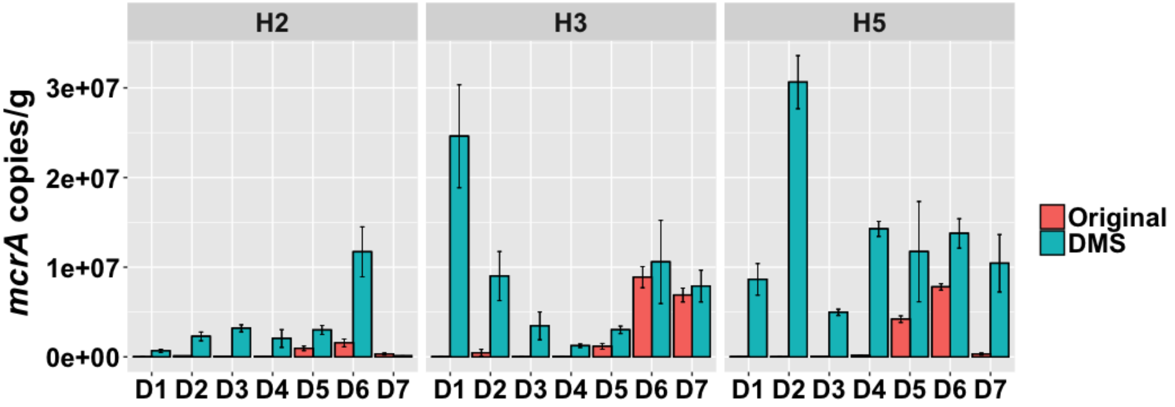


Supplementary Figure 4. Mean copy number of the *mcrA* gene per gram of wet sediment in the original and DMS-amended sediments. Error bars represent standard error above and below the average of three replicates.

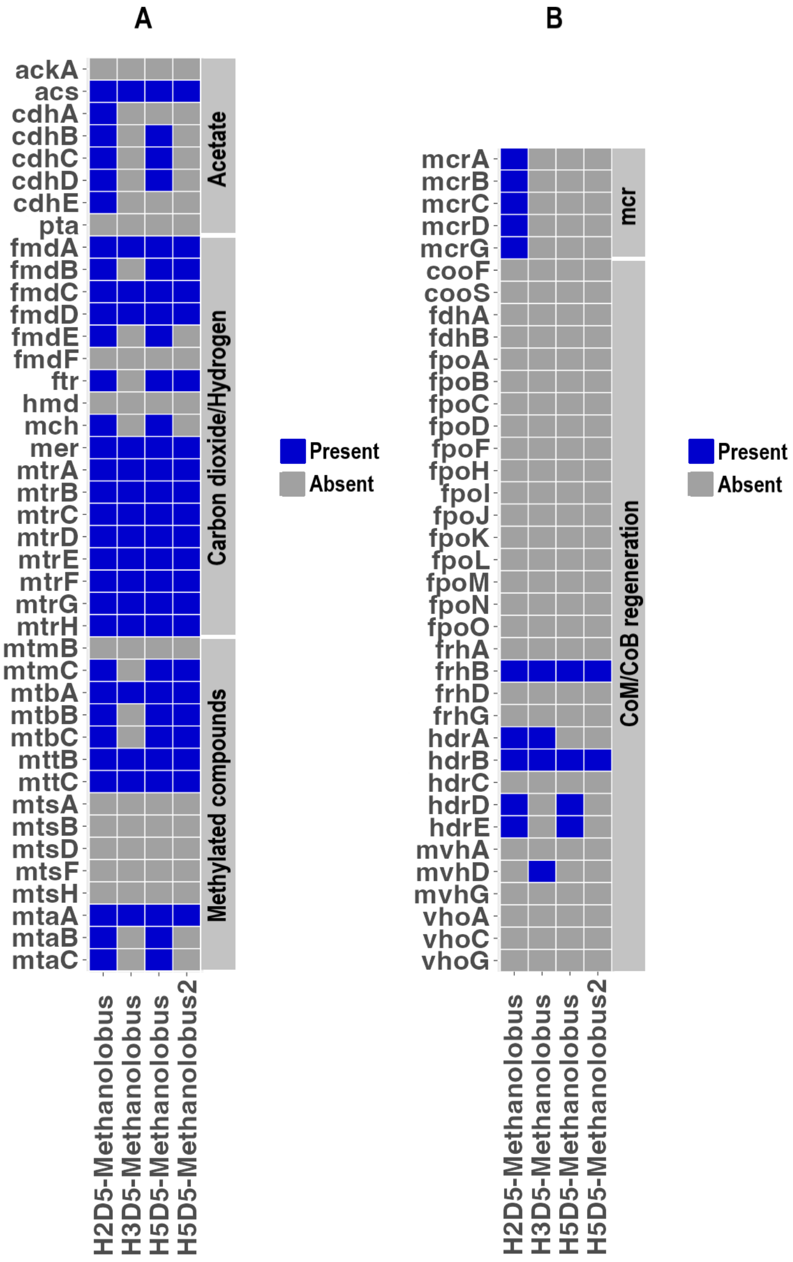


Supplementary Figure 5. Presence and absence of the genes involved in methane production in the four *Methanolobus* MAGs constructed using the metagenomics datasets. (a) Distinct genes involved in acetoclastic, hydrogenotrophic and methylotrophic methanogenesis pathways; (b) Genes common to all methanogenesis pathways.

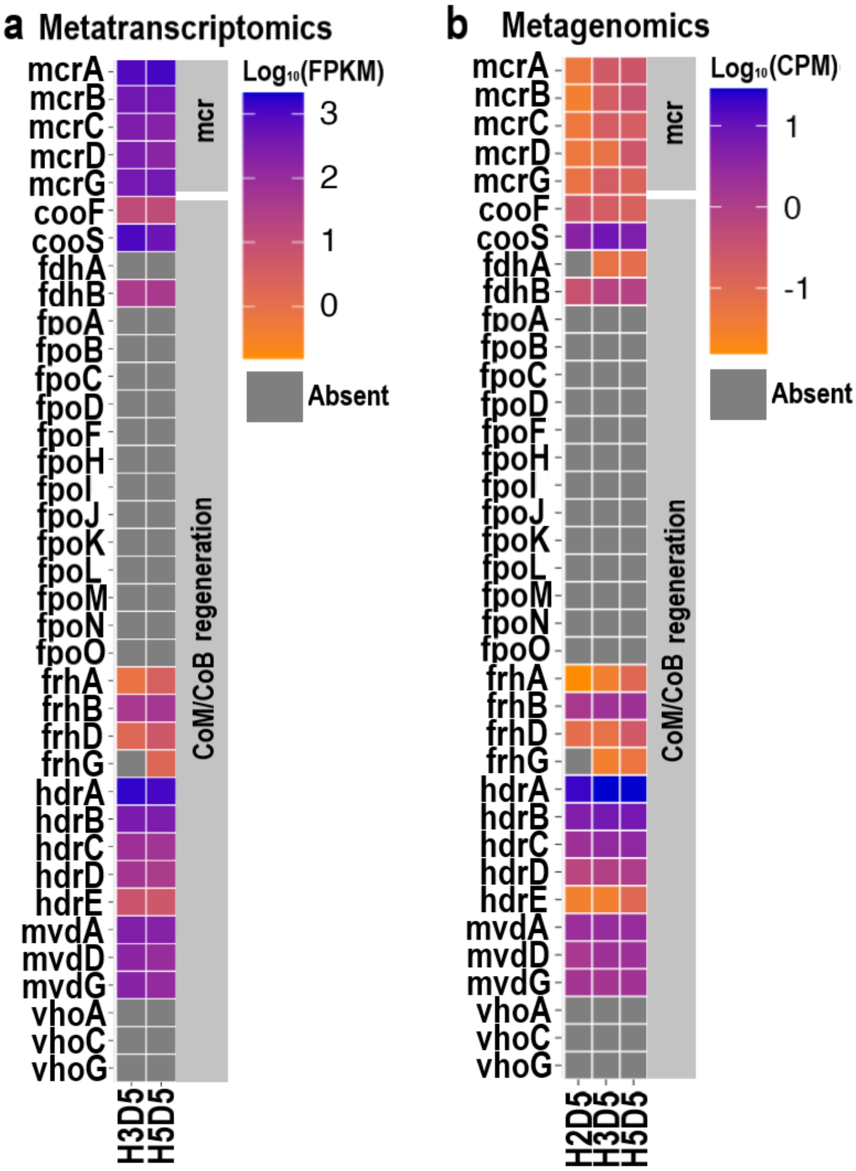


Supplementary Figure 6. Heatmap showing the normalised copy numbers of the genes common in all methanogenesis pathways. (a) Metagenomics datasets; (b) Metatranscriptomics datasets. CPM: Copies per million reads; FPKM: fragments per kilobase of gene per million reads.

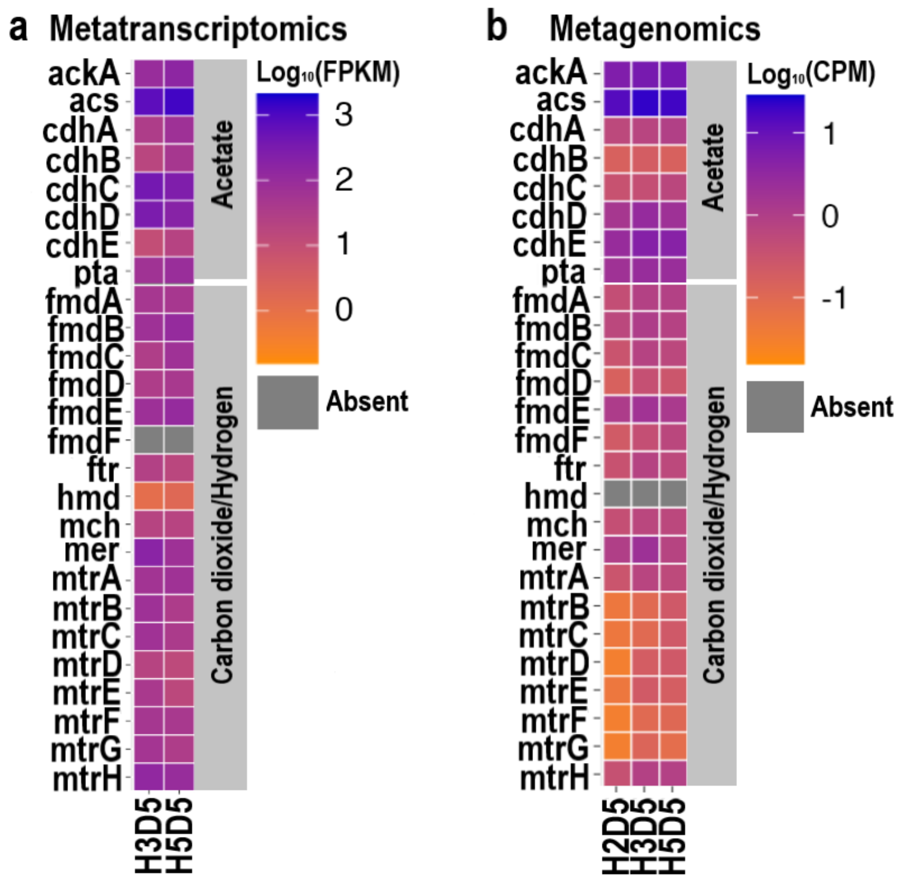


Supplementary Figure 7. Heatmap showing the normalised copy numbers of the genes involved in acetoclastic and hydrogenotrophic methanogenesis pathways. (a) Metatranscriptomics datasets; (B) Metagenomics datasets. FPKM: fragments per kilobase of gene per million reads. CPM: Copies per million reads.

Supplementary Table 1. Spearman's rank correlation coefficients ($r_s$) between total DMS consumed, total methane and $CO_2$ produced, depth, initial and end point sulfate amounts and the first two principal coordinates obtained by the *mcrA* sequence analysis. Statistically significant values are in bold. ***: p<0.001; **: p<0.01; *: p<0.05.

| **Spearman's rank correlation ($r_s$)** | **PCo1** | **PCo2** |
|---|---|---|
| DMS consumed (µmol) | **0.85*** ** | -0.17 |
| Methane produced (µmol) | **0.84*** ** | 0.13 |
| $CO_2$ produced (µmol) | **0.54*** ** | -0.03 |
| Initial sulfate (µmol) | **0.23*** | **-0.56*** ** |
| End sulfate (µmol) | **0.19*** | **-0.42*** ** |
| Depth (cm) | **-0.20*** | **0.55*** ** |

Supplementary Table 2. The list of 78 methylotrophy-related genes searched within metagenomes and metatranscriptomes.

| Genes | Metabolic Pathway | Full_Name | GeneCode |
|---|---|---|---|
| ackA | Methanogenesis (Acetate) | Acetate_kinase | K00925 |
| acs | Methanogenesis (Acetate) | Acetyl-CoA synthetase | EG11448 |
| cdhA | Methanogenesis (Acetate) | anaerobic carbon-monoxide dehydrogenase | K00192 |
| cdhB | Methanogenesis (Acetate) | anaerobic carbon-monoxide dehydrogenase | K00195 |
| cdhC | Methanogenesis (Acetate) | acetyl-CoA decarbonylase/synthase | K00193 |
| cdhD | Methanogenesis (Acetate) | acetyl-CoA decarbonylase/synthase | K00194 |
| cdhE | Methanogenesis (Acetate) | acetyl-CoA decarbonylase/synthase | K00197 |
| pta | Methanogenesis (Acetate) | phosphate acetyltransferase | K00625 |
| fmdA | Methanogenesis (CO2) | formylmethanofuran dehydrogenase | K00200 |
| fmdB | Methanogenesis (CO2) | formylmethanofuran dehydrogenase | K00201 |
| fmdC | Methanogenesis (CO2) | formylmethanofuran dehydrogenase | K00202 |
| fmdD | Methanogenesis (CO2) | formylmethanofuran dehydrogenase | K00203 |
| fmdE | Methanogenesis (CO2) | formylmethanofuran dehydrogenase | K11261 |
| fmdF | Methanogenesis (CO2) | 4Fe-4S ferredoxin | K00205 |
| ftr | Methanogenesis (CO2) | formylmethanofuran--tetrahydromethanopterin N-formyltransferase | K00672 |
| hmd | Methanogenesis (CO2) | 5,10-methenyltetrahydromethanopterin hydrogenase | K13942 |
| mch | Methanogenesis (CO2) | methenyltetrahydromethanopterin cyclohydrolase | K01499 |
| mer | Methanogenesis (CO2) | 5,10-methylenetetrahydromethanopterin reductase | K00320 |
| mtrA | Methanogenesis (CO2) | tetrahydromethanopterin S-methyltransferase subunit A | K00577 |
| mtrB | Methanogenesis (CO2) | tetrahydromethanopterin S-methyltransferase subunit B | K00578 |
| mtrC | Methanogenesis (CO2) | tetrahydromethanopterin S-methyltransferase subunit C | K00579 |
| mtrD | Methanogenesis (CO2) | tetrahydromethanopterin S-methyltransferase subunit D | K00580 |
| mtrE | Methanogenesis (CO2) | tetrahydromethanopterin S-methyltransferase subunit E | K00581 |
| mtrF | Methanogenesis (CO2) | tetrahydromethanopterin S-methyltransferase subunit F | K00582 |
| mtrG | Methanogenesis (CO2) | tetrahydromethanopterin S-methyltransferase subunit G | K00583 |
| mtrH | Methanogenesis (CO2) | tetrahydromethanopterin S-methyltransferase subunit H | K00584 |
| mtmB | Methanogenesis (Methylamine) | methylamine---corrinoid protein Co-methyltransferase | K16176 |
| mtmC | Methanogenesis (Methylamine) | monomethylamine corrinoid protein | K16177 |
| mtbA | Methanogenesis (Dimethylamine) | [methyl-Co(III) methylamine-specific corrinoid protein]:coenzyme M methyltransferase | K14082 |
| mtbB | Methanogenesis (Dimethylamine) | dimethylamine---corrinoid protein Co-methyltransferase | K16178 |
| mtbC | Methanogenesis (Trimethylamine) | dimethylamine corrinoid protein | K16179 |
| mttB | Methanogenesis (Trimethylamine) | trimethylamine---corrinoid protein Co-methyltransferase | K14083 |
| mttC | Methanogenesis (Trimethylamine) | trimethylamine corrinoid protein | K14084 |
| mtsA | Methanogenesis (Dimethylsulfide, methanethiol, methylpropionate) | methylthiol:coenzyme M methyltransferase | K16954 |
| mtsB | Methanogenesis (Dimethylsulfide, methanethiol, methylpropionate) | methylated-thiol--corrinoid protein | K16955 |
| mtsD | Methanogenesis (Dimethylsulfide, methanethiol, methylpropionate) | methyltransferase cognate corrinoid protein [ Methanosarcina acetivorans C2A ] | MA0859 |
| mtsF | Methanogenesis (Dimethylsulfide, methanethiol, methylpropionate) | cobalamin-dependent protein [ Methanosarcina acetivorans C2A ] | MA4384 |
| mtsH | Methanogenesis (Dimethylsulfide, methanethiol, methylpropionate) | cobalamin-dependent protein [ Methanosarcina acetivorans C2A ] | MA4558 |
| mtaA | Methanogenesis (Methanol) | [methyl-Co(III) methanol/glycine betaine-specific corrinoid protein]:coenzyme M methyltransferase | K14080 |
| mtaB | Methanogenesis (Methanol) | methanol---5-hydroxybenzimidazolylcobamide Co-methyltransferase | K04480 |
| mtaC | Methanogenesis (Methanol) | methanol corrinoid protein | K14081 |
| mcrA | Coenzyme M reduction to methane | methyl-coenzyme M reductase alpha subunit | K00399 |
| mcrB | Coenzyme M reduction to methane | methyl-coenzyme M reductase beta subunit | K00401 |
| mcrC | Coenzyme M reduction to methane | methyl-coenzyme M reductase subunit C | K03421 |
| mcrD | Coenzyme M reduction to methane | methyl-coenzyme M reductase subunit D | K03422 |
| mcrG | Coenzyme M reduction to methane | methyl-coenzyme M reductase subunit gamma | K00402 |
| cooF | Coenzyme B/Coenzyme M regeneration | anaerobic carbon-monoxide dehydrogenase iron sulfur subunit | K00196 |
| cooS | Coenzyme B/Coenzyme M regeneration | anaerobic carbon-monoxide dehydrogenase catalytic subunit | K00198 |
| fdhA | Coenzyme B/Coenzyme M regeneration | glutathione-independent formaldehyde dehydrogenase | K00148 |
| fdhB | Coenzyme B/Coenzyme M regeneration | formate dehydrogenase (coenzyme F420) beta subunit | K00125 |
| fpoA | Coenzyme B/Coenzyme M regeneration | F420H2 dehydrogenase subunit A | K22158 |
| fpoB | Coenzyme B/Coenzyme M regeneration | F420H2 dehydrogenase subunit B | K22159 |
| fpoC | Coenzyme B/Coenzyme M regeneration | F420H2 dehydrogenase subunit C | K22160 |
| fpoD | Coenzyme B/Coenzyme M regeneration | F420H2 dehydrogenase subunit D | K22161 |
| fpoF | Coenzyme B/Coenzyme M regeneration | F420H2 dehydrogenase subunit F | K22162 |
| fpoH | Coenzyme B/Coenzyme M regeneration | F420H2 dehydrogenase subunit H | K22163 |
| fpoI | Coenzyme B/Coenzyme M regeneration | F420H2 dehydrogenase subunit I | K22164 |
| fpoJ | Coenzyme B/Coenzyme M regeneration | F420H2 dehydrogenase subunit J | K22165 |
| fpoK | Coenzyme B/Coenzyme M regeneration | F420H2 dehydrogenase subunit K | K22166 |
| fpoL | Coenzyme B/Coenzyme M regeneration | F420H2 dehydrogenase subunit L | K22167 |
| fpoM | Coenzyme B/Coenzyme M regeneration | F420H2 dehydrogenase subunit M | K22168 |
| fpoN | Coenzyme B/Coenzyme M regeneration | F420H2 dehydrogenase subunit N | K22169 |
| fpoO | Coenzyme B/Coenzyme M regeneration | F420H2 dehydrogenase subunit O | K22170 |
| frhA | Coenzyme B/Coenzyme M regeneration | coenzyme F420 hydrogenase subunit alpha | K00440 |
| frhB | Coenzyme B/Coenzyme M regeneration | coenzyme F420 hydrogenase subunit beta | K00441 |
| frhD | Coenzyme B/Coenzyme M regeneration | coenzyme F420 hydrogenase subunit delta | K00442 |
| frhG | Coenzyme B/Coenzyme M regeneration | coenzyme F420 hydrogenase subunit gamma | K00443 |
| hdrA | Coenzyme B/Coenzyme M regeneration | heterodisulfide reductase | K03388 |
| hdrB | Coenzyme B/Coenzyme M regeneration | heterodisulfide reductase | K03389 |
| hdrC | Coenzyme B/Coenzyme M regeneration | heterodisulfide reductase | K03390 |
| hdrD | Coenzyme B/Coenzyme M regeneration | heterodisulfide reductase | K08264 |
| hdrE | Coenzyme B/Coenzyme M regeneration | heterodisulfide reductase | K08265 |
| mvdA | Coenzyme B/Coenzyme M regeneration | F420-non-reducing hydrogenase large subunit | K14126 |
| mvdD | Coenzyme B/Coenzyme M regeneration | F420-non-reducing hydrogenase iron-sulfur subunit | K14127 |
| mvdG | Coenzyme B/Coenzyme M regeneration | F420-non-reducing hydrogenase small subunit | K14128 |
| vhoA | Coenzyme B/Coenzyme M regeneration | methanophenazine hydrogenase, large subunit | K14068 |
| vhoC | Coenzyme B/Coenzyme M regeneration | methanophenazine hydrogenase, cytochrome b subunit | K14069 |
| vhoG | Coenzyme B/Coenzyme M regeneration | methanophenazine hydrogenase | K14070 |

Supplementary Table 3. Metagenome assembled genomes (MAGs) constructed from metagenome datasets from each sampling station at 19-22 cm of depth. Quality is based on the MIMAG (Bowers et al., 2017). Comp: Completeness; Cont: Contamination. Methanogen MAGs are in bold.

| Site | Kingdom | Organism | Comp | Cont | Bases | Genes | Quality |
|---|---|---|---|---|---|---|---|
| Station H2 | Bacteria | *Sideroxydans (Nitromonadales)* | 99.37% | 0.03% | 2,570,235 | 2,553 | Medium |
| | Bacteria | *Sulfuricella (Nitrosomonadales)* | 98.66% | 1.18% | 3,054,443 | 3,112 | Medium |
| | Bacteria | *Sulfurimonas (Campylobacterales)* | 98.36% | 2.12% | 2,640,566 | 2,642 | Medium |
| | Bacteria | *Sulfurivermis (Thiohalomonadales)* | 97.89% | 0.94% | 3,721,594 | 3,717 | Medium |
| | Bacteria | *Thiobacillus (Burkholderiales)* | 97.37% | 4.37% | 3,104,052 | 3,228 | Medium |
| | Bacteria | *SLDE01 (Thiohalomonadales)* | 91.02% | 2.11% | 2,870,290 | 2,863 | Medium |
| | Bacteria | *SPDF01 (Gemmatimonadales)* | 84.84% | 7.24% | 2,284,254 | 2,441 | Medium |
| | Bacteria | *Mor1 (Acidobacteriota)* | 84.22% | 4.70% | 3,172,399 | 3,142 | Medium |
| | Bacteria | *UBA2270 (Desulfobulbales)* | 83.70% | 0% | 2,242,279 | 2,255 | Medium |
| | Bacteria | *M0040 (Desulfuromonadales)* | 78.40% | 1.45% | 2,612,526 | 2,764 | Medium |
| | Bacteria | *UBA9959 (Elusimicrobiales)* | 66.07% | 1.71% | 2,008,712 | 2,008 | Medium |
| | Bacteria | *UBA2258* | 64.75% | 0.20% | 2,201,006 | 2,164 | Medium |
| | Bacteria | *GWC2-71-9 (Gemmatimonadales)* | 62.74% | 4.50% | 1,941,363 | 2,018 | Medium |
| | Bacteria | *Lutibacter (Flavobacteriales)* | 61.48% | 2.74% | 1,850,908 | 1,961 | Medium |
| | Bacteria | *Pontiella (Kiritimatiellales)* | 58.46% | 0.54% | 2,736,742 | 2,701 | Medium |
| | Bacteria | *CG2-30-66-27 (MBNT15)* | 52.66% | 0.84% | 982,015 | 1,132 | Medium |
| | Bacteria | *Ignavibacteriaceae (Ignavibacteriales)* | 52.42% | 2.33% | 1,514,737 | 1,573 | Medium |
| | Bacteria | *SMWR01 (UBA9160)* | 50.42% | 6.45% | 2,672,116 | 2,872 | Medium |
| | **Archaea** | ***Methanolobus (Methanosarcinales)*** | **92.81%** | **1.96%** | **2,489,475** | **2,669** | **Medium** |
| | Archaea | *UBA7939 (Methanosarcinales)* | 87.58% | 0.65% | 2,224,045 | 2,673 | Medium |
| Station H3 | Bacteria | *Thiobacillus (Burkholderiales)* | 100% | 0.48% | 3,269,799 | 3,337 | Medium |
| | Bacteria | *Sulfuricella (Nitrosomonadales)* | 99.29% | 0.98% | 2,882,075 | 2,947 | Medium |
| | Bacteria | *Gemmatimonadetes* | 93.20% | 4.95% | 3,005,382 | 2,914 | Medium |
| | Bacteria | *UBA9214 (Thiohalobacterales)* | 77.45% | 1.90% | 2,684,927 | 2,952 | Medium |
| | Bacteria | *Methylophagaceae (Nitrosococcales)* | 72.41% | 0.00% | 2,112,498 | 2,215 | Medium |
| | Bacteria | *UBA8639 (Nitrospirales)* | 58.53% | 4.02% | 1,749,825 | 1,957 | Medium |
| | Bacteria | *Ignavibacterium (Ignavibacteriales)* | 57.37% | 7.94% | 1,714,699 | 1,870 | Medium |
| | Bacteria | *CG2-30-66-27 (MBNT15)* | 55.57% | 0.84% | 1,184,603 | 1,340 | Medium |
| | Bacteria | *SPDF01 (Gemmatimonadales)* | 53.44% | 7.14% | 1,471,751 | 1,636 | Medium |
| | Bacteria | *BM004 (Desulfobulbales)* | 53.06% | 1.81% | 1,085,505 | 1,216 | Medium |
| | Archaea | *UBA10834 (Thermoplasmata)* | 83.37% | 1.20% | 1,375,321 | 1,451 | Medium |
| | **Archaea** | ***Methanolobus (Methanosarcinales)*** | **66.74%** | **1.31%** | **1,070,939** | **1,261** | **Medium** |
| Station H5 | Bacteria | *Methylophagaceae (Nitrosococcales)* | 99.18% | 0.88% | 2,928,202 | 2,798 | Medium |
| | Bacteria | *Sulfuricella (Nitrosomonadales)* | 95.50% | 2.84% | 3,050,167 | 3,151 | Medium |
| | Bacteria | *M0040 (Desulfuromonadales)* | 89.22% | 0.65% | 3,047,111 | 3,157 | Medium |
| | Bacteria | *Sulfurimonas (Campylobacterales)* | 79.44% | 3.88% | 1,756,086 | 1,894 | Medium |
| | Bacteria | *UBA6164 (Gracilibacteria)* | 76.99% | 2.36% | 1,161,215 | 2,152 | Medium |
| | Bacteria | *UBA9214 (Thiohalobacterales)* | 76.14% | 7.30% | 2,418,493 | 2,607 | Medium |
| | Bacteria | *BM004 (Desulfobulbales)* | 70.72% | 1.52% | 1,624,494 | 1,775 | Medium |
| | Bacteria | *Lutibacter (Flavobacteriales)* | 65.18% | 3.28% | 1,960,362 | 2,061 | Medium |
| | Bacteria | *UBA2258* | 62.76% | 1.65% | 1,130,384 | 1,198 | Medium |
| | **Archaea** | ***Methanolobus (Methanosarcinales)*** | **88.89%** | **1.31%** | **2,579,227** | **2,760** | **Medium** |
| | **Archaea** | ***Methanolobus (Methanosarcinales)*** | **62.58%** | **0%** | **1,291,140** | **1,362** | **Medium** |
| | Archaea | *SMTZ1-45 (Thorarchaeales)* | 54.35% | 0.47% | 677,230 | 810 | Medium |
